# Iron acquisition strategies of the rock-inhabiting fungus *Knufia petricola* reveal vulnerability to chelator-based mitigation

**DOI:** 10.1101/2025.11.18.689051

**Authors:** Ruben Gerrits, Julia Schumacher, Anna A. Gorbushina

## Abstract

Fungal iron acquisition has mostly been studied for pathogens living in competitive environments where securing iron from the host is critical for proliferation. How fungi from stress-rich but competition-free material surfaces handle iron uptake is, however, unknown. We studied these processes in rock-inhabiting fungi, known for their constitutive melanin-production, extremotolerant lifestyle and colonization of exposed surfaces such as solar panels and marble monuments, by choosing *Knufia petricola* to represent this group. The characterization of targeted mutants showed that *K. petricola* produces an extracellular – yet undefined – siderophore on which it relies less than on the ferroxidase-iron permease complex for reductive iron assimilation. By disrupting melanin synthesis, it was demonstrated that this pigment can reduce and adsorb iron but nevertheless does not contribute significantly to iron acquisition. Moreover, growth towards an iron source was found to be siderophore-dependent. Ferric citrate could not be directly assimilated. Finally, it was found that *K. petricola* and other rock-inhabiting fungi exhibited high sensitivity to strong iron chelators such as EDTA. These observations not only offer a potential mitigation strategy for preventing fungal colonization of subaerial materials using iron chelators, but also highlight the specialized adaptations of these fungi to low-competition environments.

**Importance:** The rock-inhabiting fungus *Knufia petricola* acquires iron primarily via reductive iron assimilation, independently of the iron-adsorbing and -reducing melanin. Its siderophore, although extracellular and allowing chemotropism towards an iron source, is not able to obtain iron from strong iron chelators such as BPS and EDTA. This sensitivity to iron sequestration opens avenues to mitigate the fungal colonization of materials such as solar panels.

## Introduction

Iron, the fourth most abundant element in Earth’s upper continental crust (1), has since the rise of atmospheric oxygen levels around 2.4 Gya (the Great Oxidation Event, GOA) started to oxidize and precipitate in the form of oxides (2–5). Imlay (6) noted that the abundant amounts of available iron pre-GOA resulted in the use of iron-sulphur clusters within many proteins such as redox enzymes involved in respiration. Post-GOA, oxygen-enduring organisms started to thrive in aerobic environments by adopting strategies to cope with low iron availability and oxidative stress (6, 7).

Fungi may use up to four mechanisms to acquire iron from their environment (8–10): (i) uptake of ferric iron after its extracellular reduction by ferric reductases (FREs) and re-oxidation via a high-affinity ferroxidase-iron permease complex (reductive iron assimilation, RIA), (ii) uptake of ferric iron complexed by low-molecular-mass chelators called siderophores via high-affinity siderophore transporters (siderophore-mediated iron acquisition, SIA), (iii) uptake of heme-complexed iron, and (iv) uptake of ferrous iron by low-affinity transporters. Iron uptake is well studied in the budding yeast *Saccharomyces cerevisiae* and the opportunistic animal pathogen *Aspergillus fumigatus. S. cerevisiae* possesses one RIA complex localized in the plasma membrane (FTR1-FET3) (11), and one low-affinity ferrous iron transporter (FET4) (12), and although it does not synthesize siderophores, it has four siderophore iron transporters (SITs) for uptake of foreign siderophores (xenosiderophores) (13, 14). The filamentous fungus *A. fumigatus* contains one RIA complex (FtrA-FetC) (15), a low-affinity iron transporter (FetD) (10), and five putative SITs for taking up iron bound to (xeno)siderophores (16). Four hydroxamate siderophores are produced by non-ribosomal peptide synthetases (NRPSs): ferricrocin, hydroxyferricrocin, fusarinine C and triacetylfusarinine C (17). Ferricrocin was initially assumed to function solely in intracellular iron handing (10, 17), later studies showed its secretion and role in iron acquisition (18).

The mechanisms of iron acquisition, essential for growth and proliferation, have been primarily studied in pathogenic fungi to develop antifungal strategies in a medicinal or agricultural context (8, 10, 19–21). The active chelation of iron by the animal host hampers microbial infections (so-called nutritional immunity (22)), rendering the habitat of pathogenic fungi more iron depleted than that for other fungi. We hypothesize, therefore, that the absence of competition for iron in their environmental niches renders saprobic fungi more sensitive to strong iron chelators, thereby offering a basis for potential mitigation strategies. We focused on the polyphyletic group of black fungi, which share similar morpho-physiological adaptations allowing them to survive in harsh environments, including melanized cell walls, slow yeast-like or meristematic growth and the absence of complex reproduction cycles (23). While several black fungi are (opportunistic) animal pathogens (24), the group of rock-inhabiting fungi colonize sub-aerial environments such as bare rocks in cold and hot deserts and human-made structures such as marble monuments and solar panels (25–27). The increasing use of the latter surfaces for energy generation (28) together with the considerable challenge in removing black fungal colonies or mitigating their growth (29), underscores the need of novel mitigation strategies.

Despite the high diversity, only few genomes of rock-inhabiting fungi within the Arthoniomycetes, Eurotiomycetes and Dothideomycetes have been sequenced to date. For even fewer fungi, genetic engineering tools for functional analyses are available. To address this gap, we selected *Knufia petricola* (Eurotiomycetes, Chaetothyriales, Trichomeriaceae) as a model organism for related rock-inhabiting fungi, and developed tools for comparative phenotyping and genetic approaches (30–32). *K. petricola* constitutively produce the black 1,8-dihydroxynaphthalene (DHN) melanin and reddish carotenoids, which become visible in melanin-deficient mutants (30). Whether melanin, a multifunctional pigment, plays are in iron acquisition, however, remains unknown.

Here, we combined bioinformatics and genetics analyses to reveal the basis of iron acquisition in *K. petricola*. Genes for two high-affinity iron uptake systems and DHN melanin synthesis were deleted to study the capacity of the systems to acquire iron from media or minerals. Accordingly, *K. petricola* primarily employs RIA over SIA, produces an (extracellular) siderophore which enables growth towards an iron source, and relies minimally on melanin, which plays no direct role in iron acquisition. The observation that *K. petricola* and other rock-inhabiting black fungi fail to grow in presence of strong iron chelators demonstrates that iron chelation can serve as an effective and novel strategy to mitigate fungal colonization of exposed surfaces.

## Results

### Genome analysis reveals genetic potential for RIA and SIA in K. petricola

Genes, putatively involved in iron acquisition in *K. petricola* and related black fungi (Chaetothyriales), were identified based on sequence similarity to characterized proteins from *S. cerevisiae* and *A. fumigatus*. Their distribution differs markedly between pathogenic Herpotrichiellaceae and predominantly saprobic Trichomeriaceae species (Figure 1, Table S1). Most of the species lack a transporter for low-affinity iron uptake (FetD), but the number of ferroxidase–permease (FF) pairs correlates strongly with the lifestyle: pathogenic species such as *Fonsecaea predosoi* typically harbor at least two copies, whereas saprobic species harbor single copies. FREs are generally more abundant in Herpotrichiellaceae. All species have at least one ortholog of *A. fumigatus* SidJ (EST), involved in intracellular iron release from siderophores, whereas SITs are markedly more abundant in Herpotrichiellaceae species. With respect to siderophore biosynthesis, all species harbor a SidC-like NRPS, but SidD-like NRPSs and the monooxygenase SidA are absent from several Trichomeriaceae species, indicating incomplete hydroxamate siderophore pathways. Further, Herpotrichiellaceae species harbor at least two putative vacuolar iron transporters (CCC), which are missing in most Trichomeriaceae. Interestingly, this absence correlates with the presence of a putative iron-binding ferritin (FER).

**Figure 1:**
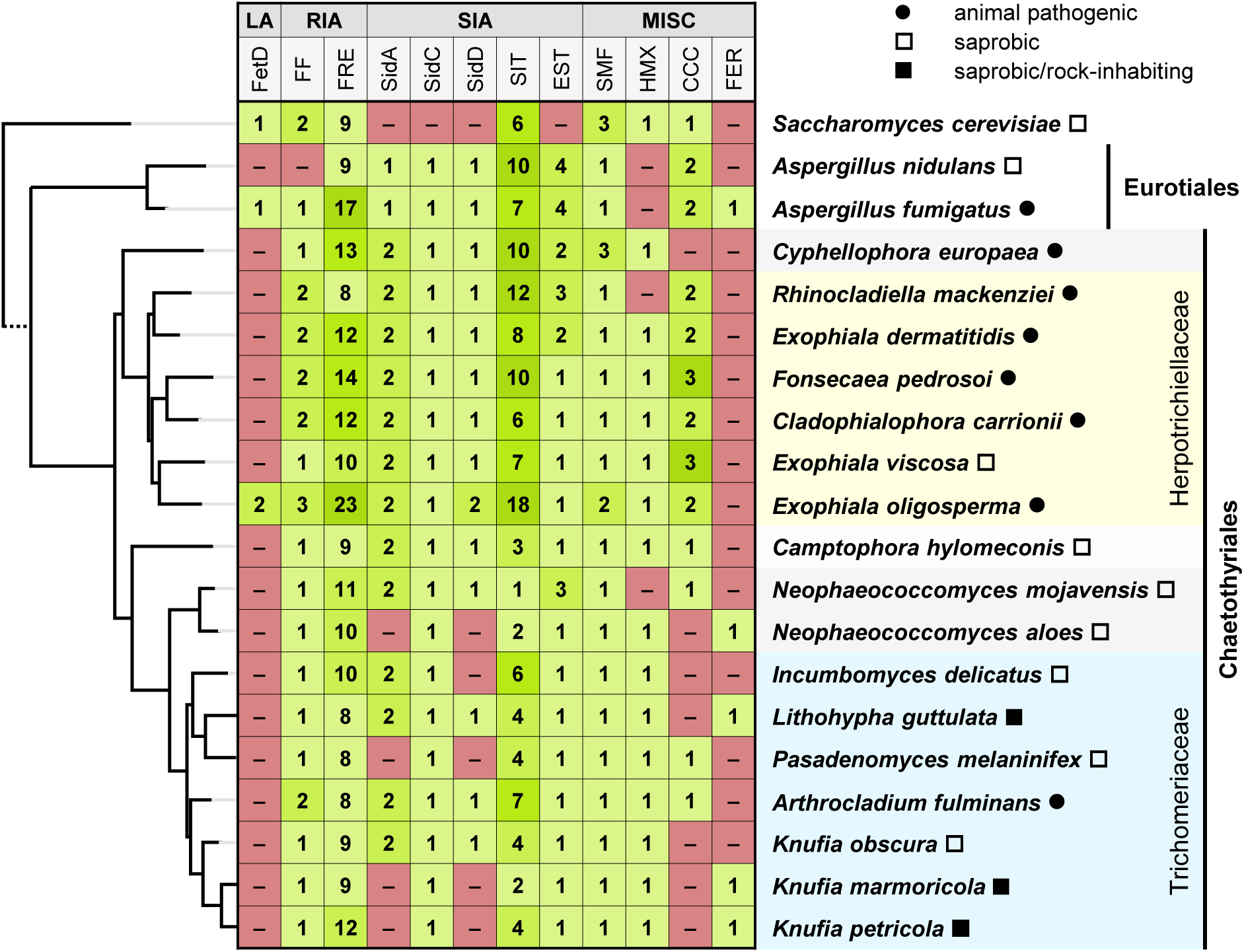
Distribution of iron acquisition genes in black fungi of the order Chaetothyriales. Representative Chaetothyriales species were selected based on phylogenetic diversity and isolation source. Species were categorized as pathogenic (repeatedly isolated from human infections), saprobic/rock-inhabiting (repeatedly isolated from rock surfaces), or saprobic (all others, for example from plant leaves, cleanroom surfaces, soil crust, ant nests). The phylogenetic tree (on the left) was constructed from concatenated protein sequences of ACT1 (actin), ADE2 (phosphoribosylaminoimidazole carboxylase), H2B (histone 2B), TEF1 (translation elongation factor 1-α), and URA3 (orotidine 5′-phosphate decarboxylase). Numbers indicate predicted iron acquisition genes identified by sequence similarity, compared to *S. cerevisiae*, *A. fumigatus*, and *A. nidulans*. Gene categories – LA (low-affinity uptake): iron transporters (FetD); RIA (reductive iron assimilation, high-affinity): ferroxidase-iron permease pairs (FF), ferric reductases (FRE); SIA (siderophore-mediated iron acquisition, high-affinity): biosynthetic enzymes (SidA, SidC, SidD), siderophore iron transporters (SIT), intracellular siderophore-hydrolyzing esterases (EST); MISC (miscellaneous): divalent metal transporters (SMF), heme oxygenases (HMX), vacuolar iron transporters (CCC), ferritins (FER). For species details, genome sequences, and protein sequences, see Table S1; for *K. petricola* genes, see also Table S2.

*K. petricola* possesses a single RIA complex (Figure S1) and twelve putative FREs (Figure S2) – more than other Trichomeriaceae, including *K. marmoricola*. Further, it has four SITs, with SIT1 clustering with *S. cerevisiae* SITs and *A. fumigatus* Sit1, while the others show varying similarity to *A. fumigatus* SITs (Figure S3). SidA- and SidD-like proteins are absent in both rock-inhabiting *Knufia* species (Figure S4), a pattern shared with *Neophaeococcomyces aloes* and *Pasadenomyces melaninifex*, which were isolated from cleanrooms and whose natural habitats remain unknown. Similarly, the absence of vacuolar iron transporters combined with the presence of a putative ferritin (Figure S5) is a feature shared by the three recognized rock-inhabiting species (*K. petricola*, *K. marmoricola*, *Lithohypha guttulata*), and *N. aloes*. Additionally, the *K. petricola* genome comprises the genes encoding the conserved transcriptional regulators SRE1, HAPX, and SRB1 (Table S2).

In summary, *K. petricola* contains single-copy genes for RIA and SIA, indicating a potentially limited iron acquisition capacity. To assess the roles of the two systems under varying iron conditions, mutants were generated and characterized (Table S3).

### RIA **–** mediated by FTR1 and FET1 **–** facilitates growth under iron limitations

The genes encoding a high-affinity iron permease (FTR1) and a ferroxidase (FET1) as components of the RIA complex, are physically linked in the genome of *K. petricola* by sharing a bidirectional promoter (Figure S1). This genomic organization is typical for fungal genomes where one to three copies of the *ftr1-fet1* cluster are commonly found. *K. petricola* FTR1 and FET1 share 61 % and 52 % aa identity, respectively, with their orthologs from *A. fumigatus*, and 69 % and 59 % identity, and 51 % and 45 % identity, with those from *E. dermatitidis*.

To investigate the role of RIA in *K. petricola*, the genomic region encompassing both genes was deleted by replacing it by a resistance cassette resulting in the mutant Δ*ftr1-fet1* (Figure S6a). Growth of the generated mutants were assayed on minimal medium (MM) under three iron conditions: iron limitation (−Fe, no added iron), iron sufficiency (+Fe, 30 µM Fe), and iron repletion (hFe, 1,511 µM Fe). When cell suspensions were dropped onto agar, the Δ*ftr1-fet1* mutant exhibited considerably reduced growth on −Fe and +Fe compared to the WT, but not under hFe conditions (Figure 2a). The quantification of the iron content in the biomass obtained from iron-limiting conditions (−Fe) yielded, however, similar results for the mutant and the WT (Figure 2b). The growth parameters of single colonies on the three solid media were quantified using the ScanLag system, which performs time-lapse scanning of Petri dishes (Figure 2c). The WT exhibited comparable growth rates across all tested conditions (approximately 0.006 mm^2^ h^-1^). However, its lag phase was significantly prolonged on hFe (148 ± 14 h) compared to −Fe (109 ± 1 h, *p* = 0.0001) and +Fe (111 ± 2 h, *p* = 0.0013). The Δ*ftr1-fet1* mutant did not grow sufficiently without iron to allow quantification of growth parameters. On +Fe, its growth rate was similar to that of the WT, but its lag phase was significantly extended (495 ± 22 h, *p* < 0.0001).

**Figure 2:**
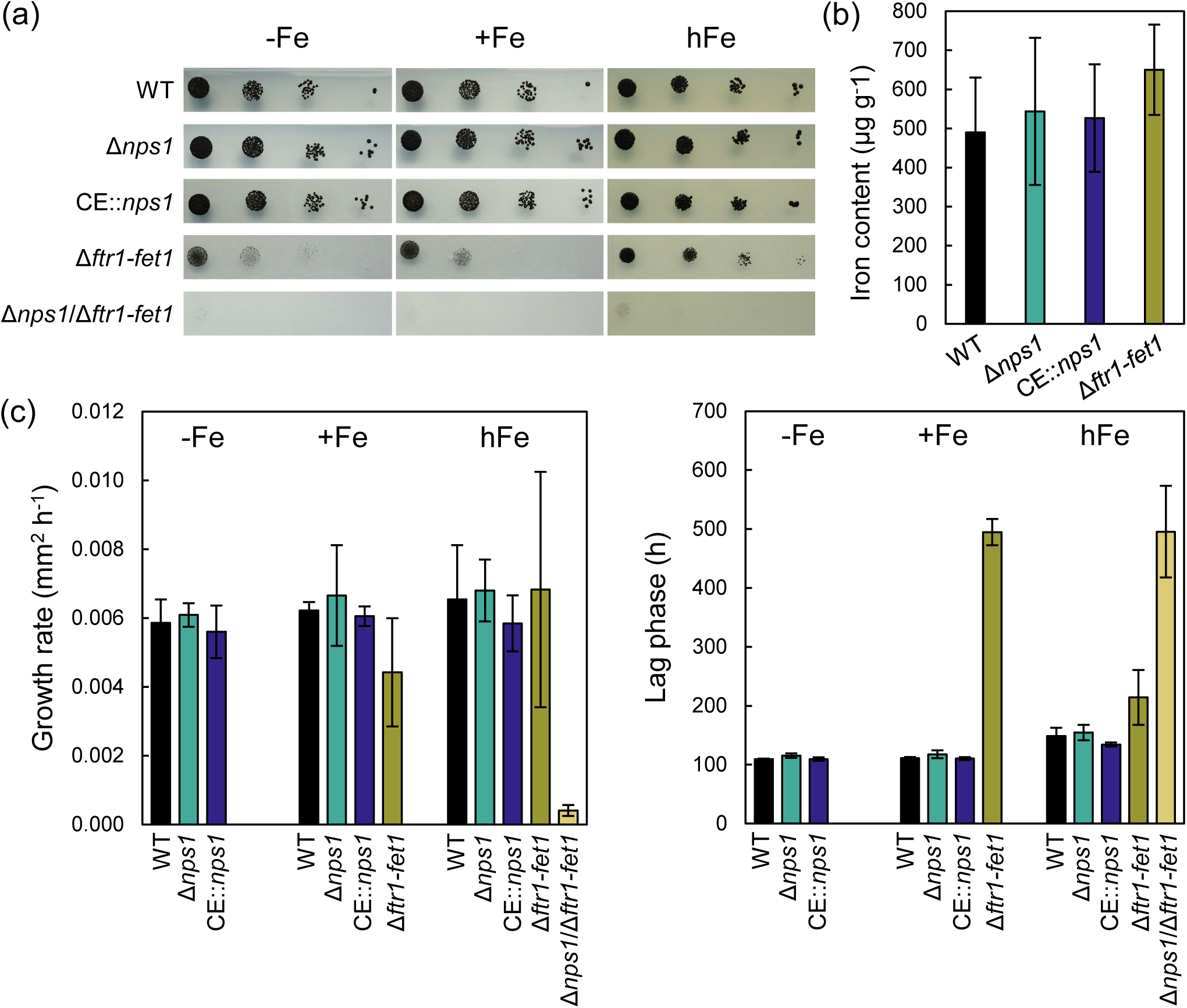
*K. petricola* acquires iron primarily through reductive iron assimilation (RIA) by FTR1–FET1. (a) RIA-deficient Δ*ftr1-fet1* mutants fail to grow under iron-limited conditions. Ten-microliter droplets containing 10^4^, 10^3^, 10^2^, and 10^1^ colony forming units (CFU) of selected strains were spotted onto solid medium without iron (–Fe), or supplemented with 30 µM iron (+Fe) or 1,511 µM iron (hFe) and incubated for 12 days. (b) WT and iron uptake-deficient mutants accumulate comparable amounts of iron in their biomass. The iron content was quantified for biomass harvested from 21-d-old cultures grown on solid +Fe medium. The biomass was digested with 1 M HNO3, and analyzed by inductively coupled plasma-optical emission spectroscopy (ICP-OES). Shown are the averages of three independent replicates with two times the standard error or analytical uncertainty, whichever was highest. (c) High-iron conditions improve the growth of RIA-deficient mutants, but not to wild-type levels. Growth rates and lag phases on media with different iron contents were recorded by the ScanLag system. Shown are the averages of three independent replicates with two times the standard error.

To verify that the observed growth phenotype of the Δ*ftr1-fet1* mutant was specifically due to the deletion of the *ftr1-fet1* locus, a complementation strain was generated by reinserting the *ftr1-fet1* pair into the neutral genomic site *igr2* (Figure S6b). The resulting strain, Δ*ftr1-fet1*::*ftr1-fet1*, exhibited growth parameters comparable to those of the WT, including similar growth rates and lag phases under both iron-limiting and iron-sufficient conditions (Figure S7). These results demonstrate that the growth defect of the mutant is due to the loss of *ftr1* and *fet1*, confirming their role in RIA. However, the observation that the mutant retains limited growth under low-iron conditions (−Fe and +Fe) suggests the presence of an alternative iron uptake system that partially compensates for the loss of *ftr1*-*fet1*.

### SIA – mediated by an extracellular, yet unidentified siderophore – requires NPS1

NPS1, comprising 4,897 aa, represents the sole NRPS of *K. petricola*. It shares 22 % and 41 % aa identity with SidC of *A. fumigatus* and *F. pedrosoi*, respectively, both known to participate in ferrocrocin biosynthesis (17, 33). Both Chaetothyriales NRPSs feature an additional tandem of thiolation (T) and condensation (C) domains (Figure S4). However, ferricrocin could not be detected in either the culture supernatant or biomass of *K. petricola* (data not shown), consistent with the absence of a *sidA* ortholog (Figure 1) – a gene needed for the production of hydroxamate siderophores such as ferricrocin (8) and typically located upstream of the NRPS-encoding gene.

To investigate whether *nps1* is involved in iron acquisition, the gene was deleted (Δ*nps1*) and its expression was elevated (Figure S8). For the latter, the native promoter of *nps1* was replaced with the strong P*oliC* from *A. nidulans*, resulting in the strain CE::*nps1* (constitutive expression of *nps1*). To explore a potential functional interdependency between the RIA system and NPS1, the *ftr1-fet1* locus was additionally deleted in the Δ*nps1* background, generating the double mutant Δ*nps1*/Δ*ftr1-fet1*. Neither the deletion nor the constitutive expression of *nps1* resulted in any visible or quantifiable changes of colony growth under any of the three iron conditions tested. This included comparable iron contents in the biomass under iron-limited conditions (Figure 2). In contrast, the Δ*nps1*/Δ*ftr1-fet1* mutant failed to grow on −Fe and +Fe, indicating a synthetic growth defect under low-iron conditions. On hFe, growth was partially restored, although the strain exhibited a significantly prolonged lag phase (495 ± 78 h, p = 0.017) and a reduced growth rate (0.0004 ± 0.0002 mm^2^ h^-1^, *p* = 0.252) compared to the WT.

Given that NPS1 appears to contribute to iron acquisition and becomes essential in the absence of a functional RIA system, we next investigated whether the NRPS might be involved in the production of a diffusible, iron-binding compound, potentially a previously unidentified siderophore. A cross-feeding assay was performed, in which the Δ*nps1*/Δ*ftr1-fet1* mutant was grown under iron-limiting conditions in proximity to either the WT, Δ*nps1*, Δ*ftr1-fet1*, or CE::*nps1*. This allowed to assess whether any of these strains could secrete an iron-chelating compound capable of rescuing the growth of the double mutant. In addition, the iron-lacking (−Fe) medium was supplemented with either iron-free forsterite (Mg_2_SiO_4_) or iron-containing olivine ((Mg,Fe)_2_SiO_4_) to evaluate whether mineral-derived iron could support growth under these conditions (Figure 3a). Growth of the Δ*nps1*/Δ*ftr1-fet1* mutant was absent when co-cultivated with Δ*nps1*, but detectable next to the WT and most pronounced in proximity to Δ*ftr1-fet1* and CE::*nps1* colonies. Interestingly, similar growth of the double mutant was observed for media without and with iron-free forsterite, whereas growth was reduced in the presence of the iron-containing olivine. Together, these observations indicate that NPS1 is involved in the production of an extracellular, diffusible siderophore. The reduced growth stimulation in the presence of olivine suggests lower siderophore production. In an alternative experiment, olivine and forsterite were used to investigate whether WT, Δ*nps1*, and Δ*ftr1-fet1* can actively grow toward an iron source, as an indication of chemotropism (Figure 3b). The strains were inoculated between the two minerals on −Fe medium lacking a nitrogen source to promote filamentous growth. This revealed, that the Δ*nps1* mutant did show significantly (*p* = 0.280) reduced growth towards the olivine than the WT. The minimal filamentous growth of Δ*ftr1-fet1* mutant prevented the quantification of its chemotropism. These results imply that the unknown siderophore produced by *K. petricola* likely facilitates iron acquisition from olivine.

**Figure 3:**
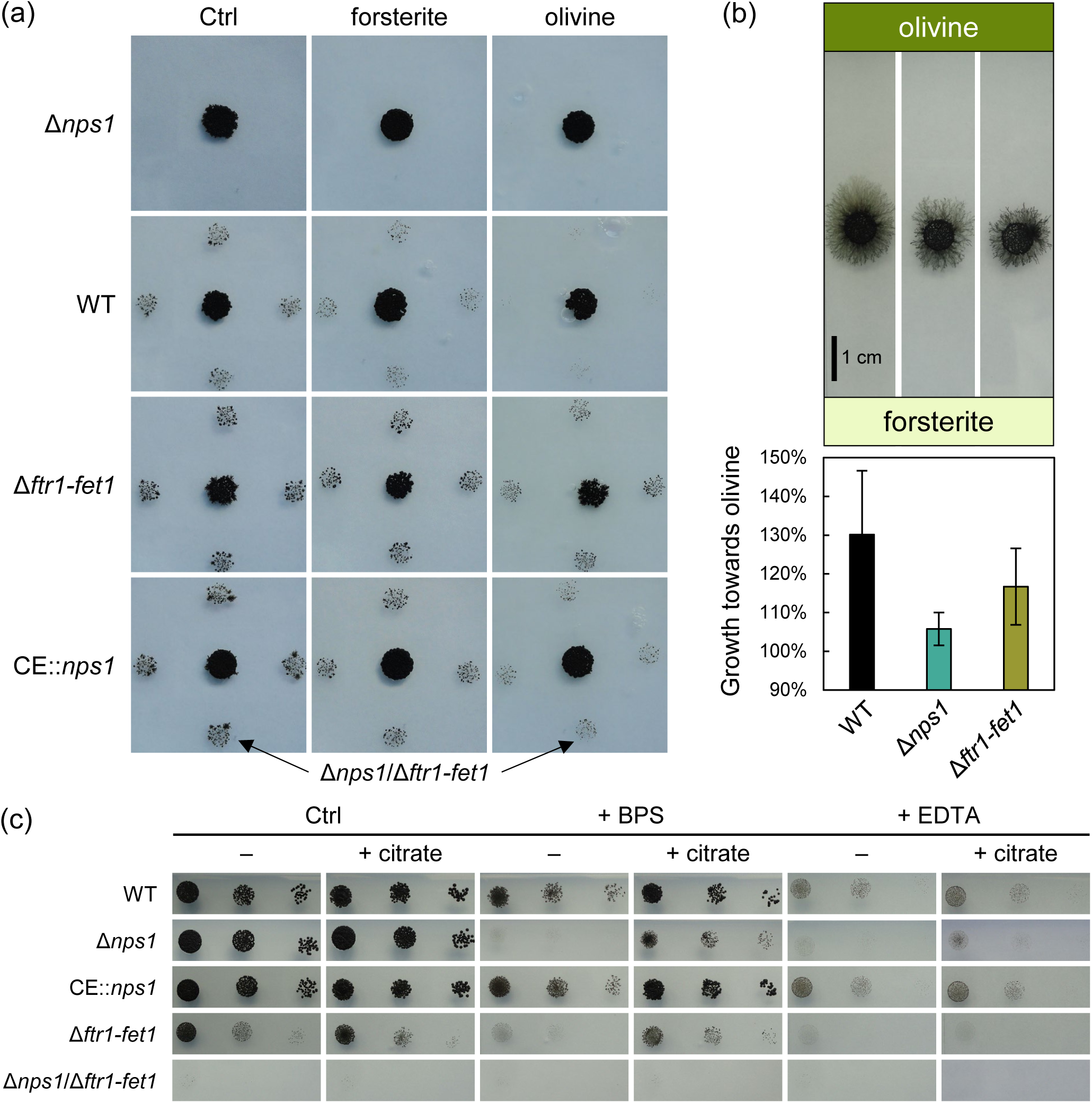
NPS1 contributes to siderophore-mediated iron acquisition (SIA) in *K. petricola*. (a) NPS1 is required to produce an extracellular siderophore. Four 10 µl drops each containing 10^3^ CFU of the Δ*nps1*/Δ*ftr1-fet1* mutant were spotted next to drops of either Δ*nps1*, WT, Δ*ftr1-fet1* or CE::*nps1* on solid –Fe medium alone (Ctrl) or supplemented with forsterite (Mg2SiO4) or olivine ((Mg,Fe)2SiO4). Growth of Δ*nps1*/Δ*ftr1-fet1* indicates production of a diffusible siderophore by the neighboring strain after 34 days of incubation. (b) The extracellular siderophore enables chemotropism towards an iron source. Five 10 µl drops containing 10^3^ CFU of WT, Δ*nps1* or Δ*ftr1-fet1* were spotted onto iron-lacking agar in between a streak of iron-rich olivine and iron-free forsterite. The relative growth rates towards olivine (growth towards olivine/growth towards forsterite, %) were calculated after 74 days of incubation. Given are the averages of three independent replicates (with five colonies each) with two times the standard error. (c) NPS1 is not required for growth on iron-deficient medium when RIA is operating. 10^4^, 10^3^, and 10^2^ CFU of selected strains were dropped onto iron-lacking (–Fe) agar with and without 1 mM citrate, 4.5 µM BPS or 4.0 µM EDTA and incubated for 12 days.

To confirm the growth phenotypes of the Δ*nps1* mutant, a complementation strain was constructed by reinserting *nps1* into its native locus through the *in vivo* assembly of the large coding region from four overlapping amplicons (Figure S8). Restoration of siderophore production in the Δ*nps1*::*nps1* strain was demonstrated by its ability to support growth of the adjacent Δ*nps1*/Δ*ftr1-fet1* mutant (Figure S9). In sum, the data demonstrate that a SIA system operates in *K. petricola*, involving an extracellular, yet unidentified siderophore.

### K. petricola and other rock inhabitants exhibit high sensitivity to iron depletion

The addition of chelators to the culture medium intensifies iron limitation by binding residual trace iron, even in media considered iron-free. Under such controlled conditions, both the iron-binding strength of metabolites, such as siderophores, and the total iron-binding strength of organisms can be experimentally assessed and compared.

When iron-lacking medium (−Fe) was supplemented with 4.5 µM of the ferrous iron chelator bathophenanthroline disulfonate (BPS) or 4.0 µM of the ferric iron chelator ethylenediaminetetraacetic acid (EDTA) (Figure 3c), all tested strains exhibited reduced growth: WT and CE::*nps1* showed comparable reduced growth, while growth of both Δ*nps1* and Δ*ftr1-fet1* was nearly abolished. Notably, the growth inhibition of Δ*nps1* was more pronounced, considering that this mutant displayed more robust growth on −Fe medium without chelators compared to Δ*ftr1-fet1*. The complemented strain Δ*nps1::nps1* exhibited BPS sensitivity comparable to that of the WT (Figure S9). Interestingly, supplementation with 1 mM citrate enabled the Δ*nps1* mutant to grow in the presence of BPS and EDTA to nearly the same extent as the WT (Figure 3c), suggesting that citrate may partially compensate for the loss of siderophore function. The Δ*ftr1-fet1* mutant also showed enhanced growth upon citrate addition. However, no improvement was observed for the double mutant Δ*nps1*/Δ*ftr1-fet1*, indicating that both iron acquisition pathways are required for citrate-supported growth under chelator stress.

To assess whether other rock-inhabiting black fungi are similarly sensitive to iron chelators as *K. petricola*, three species isolated from solar panels and belonging to different classes were tested: BAM-BF001 (Eurotiomycetes), BAM-BF027 (Arthoniomycetes), and BAM-BF046 (Dothideomycetes). As a reference, *S. cerevisiae* was included, which is known to thrive under iron-limited conditions (34). The minimal inhibitory concentrations (MICs) of the two iron chelators were determined by inoculating malt extract agar (MEA) with varying BPS and EDTA concentrations with CFU suspensions and quantifying colony formation after two weeks of incubation. All rock-inhabiting black fungi exhibited comparable sensitivities to these chelators with MICs of 150 µM for BPS and 200-250 µM for EDTA. By contrast, *S. cerevisiae* was less sensitive with MICs of 500 µM for BPS and 750 µM for EDTA (Figure 4). Taking together, iron depletion by adding iron chelators to the medium (further) reduced growth of RIA- and SIA-deficient *K. petricola* mutants suggesting that the NPS1-derived siderophore is a weak one. This growth inhibition could be partially reversed by adding citrate, demonstrating that iron-citrate is an iron source. Moreover, *K. petricola* shares the sensitivity to iron chelators with other unrelated rock-inhabiting black fungi.

**Figure 4:**
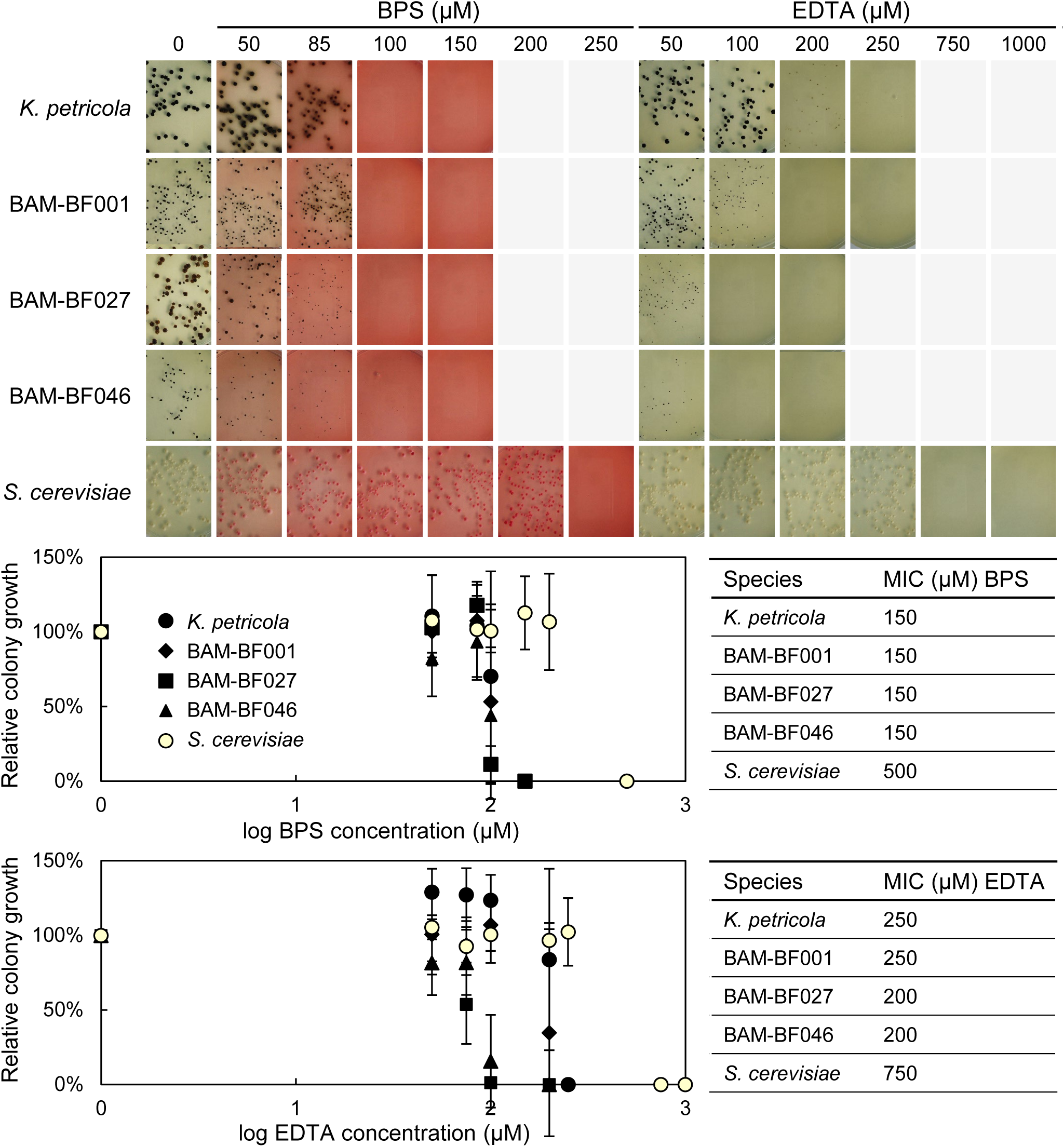
*K. petricola* and other rock-inhabiting black fungi are more sensitive towards iron chelators than *S. cerevisiae*. 200 to 450 CFU of *K. petricola* WT (A95), rock-inhabiting fungi (BAM-BF001, BAM-BF027 and BAM-BF046) and *S. cerevisiae* DSM 1333 were distributed on MEA, that was buffered at pH 6 and containing BPS or EDTA in varying concentrations. After 14 days of incubation, images were taken, and the visible colonies were counted for calculating relative colony growth in percent [number of colonies on supplemented MEA/number of colonies on MEA without chelator × 100] of each replicate. Shown are the averages of three independent replicates with two times the standard error. The minimum inhibitory concentration (MIC) of BPS and EDTA was defined as the lowest concentration at which the relative colony growth was 0 %.

### Melanin adsorbs and reduces iron but is not involved in iron acquisition

To investigate the role of melanin and its metal-binding properties in iron acquisition, melanin-deficient RIA and SIA mutants were generated. For this, *pks1*, encoding the polyketide synthase essential for DHN melanin synthesis, was deleted in Δ*nps1* and Δ*ftr1-fet1* backgrounds (Figure S10) and compared with the WT and the Δ*pks1* mutant (30).

In a drop assay, the sensitivity of the strains to oxidative stress induced by hydrogen peroxide (H_2_O_2_) under different iron conditions was evaluated. All strains showed reduced growth with H_2_O_2_: Δ*nps1* and CE::*nps1* were equally sensitive to H_2_O_2_ as the WT, whereas all other strains failed to grow with H_2_O_2_ highlighting that Δ*pks1* and Δ*pks1*/Δ*nps1* mutants as more sensitive to H_2_O_2_ compared to their melanized counterparts (WT, Δ*nps1*) (Figure 5a). The dark pigmentation and the tolerance to H_2_O_2_ was found restored in the control strain Δ*pks1*::*pks1*, generated in this study by introducing *pks1* with its native regulatory regions into the *igr2* locus of Δ*pks1* (Figure S10).

**Figure 5:**
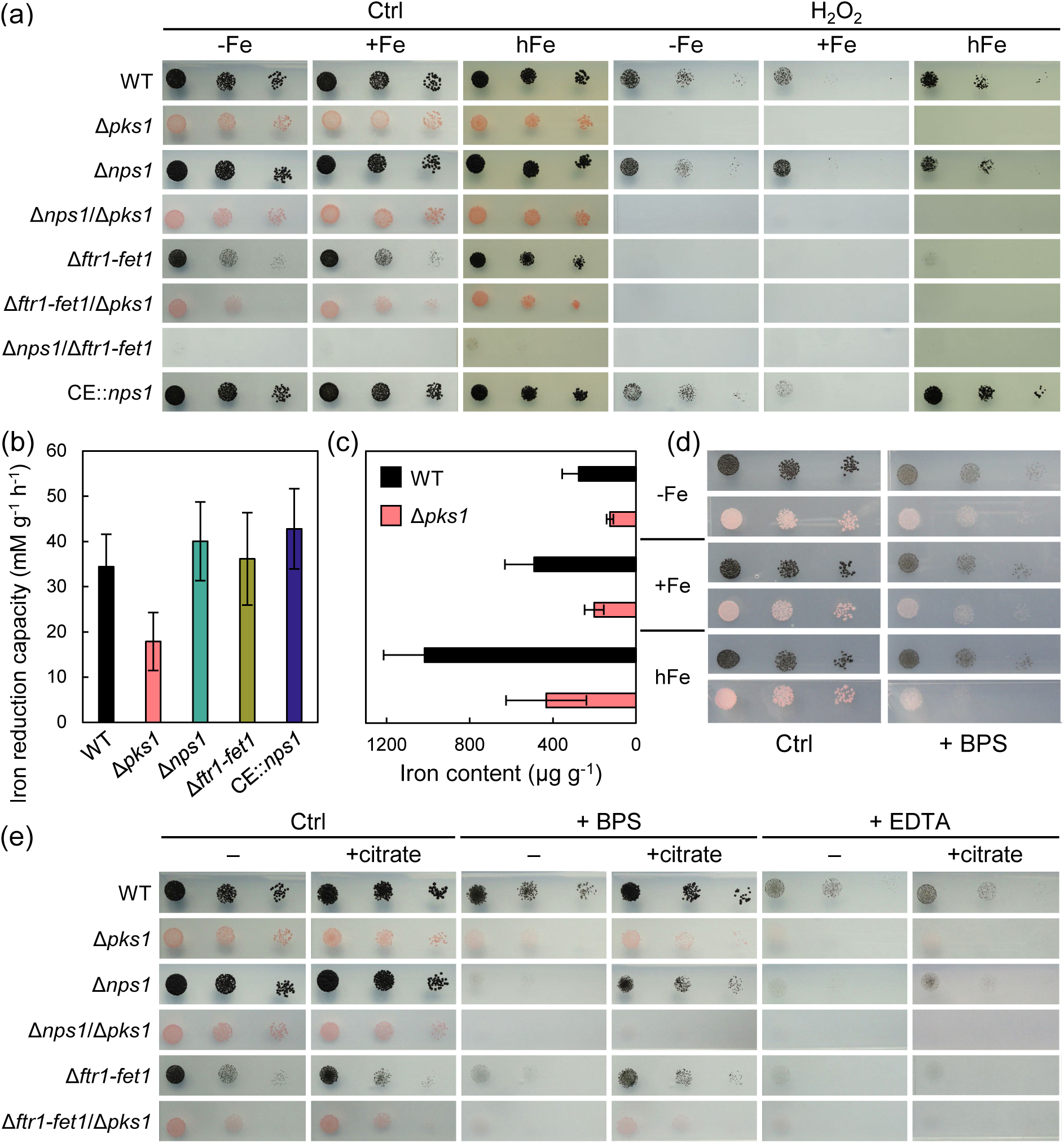
The melanin of *K. petricola* facilitates iron reduction and adsorption but is not involved in iron uptake. (a) Non-melanized mutants exhibit increased sensitivity to oxidative stress induced by hydrogen peroxide (H2O2) – independent of the iron content. CFU of melanized and non-melanized Δ*pks1* strains were point-inoculated on –Fe, +Fe and hFe agar without and with 2 mM (–Fe and +Fe) or 2.75 mM (hFe) H2O2, and incubated for 12 days. (b) The non-melanized Δ*pks1* mutant, in contrast to RIA- and SIA-deficient strains, exhibits attenuated iron-reducing activity. Liquid –Fe medium was inoculated with 10^6^ CFU of the indicated strains and incubated for 14 days. The iron reduction capacity – expressed as mM Fe(II) g^-1^ biomass h^-1^ – was determined by measuring the absorbance at 535 nm (A535) of the cultures after incubation with the iron-chelator BPS. Shown are the averages of four independent replicates with two times the standard error. (c) The biomass of the non-melanized Δ*pks1* mutant contains less iron compared to that of the WT. The two strains were cultivated for 21 days on solid –Fe, +Fe and hFe medium. Iron contents – expressed as µg iron per g biomass – were quantified via ICP-OES analyses of the digested biomass. Shown are the averages of three independent replicates with two times the standard error or analytical uncertainty, whichever was highest. (d) The melanized WT (black) and the non-melanized Δ*pks1* mutant (pink) exhibit similar growth on iron-depleted conditions, regardless of the iron availability during precultivation. The two strains were cultivated for seven days on agar without iron (–Fe) or with iron (+Fe, or hFe). Cells from these cultures were then spotted onto –Fe agar supplemented with 4.5 µM BPS as indicated. Images were taken after 12 days of incubation. (e) The non-melanized mutants were equally sensitive towards strong iron chelators than the melanized strains. Indicated strains were point-inoculated on solid MM medium without iron (–Fe) and with or without 1 mM citrate, 4.5 µM BPS and/or 4.0 µM EDTA. Results were documented after 12 days of incubation.

The iron-reducing capacity was assessed in both the culture supernatants and biomass of WT, Δ*pks1,* and the melanized RIA- and SIA-deficient mutants. The biomass of the Δ*pks1* mutant exhibited a significantly lower iron-reducing capacity compared to the other strains (e.g., 53 % of WT, *p* = 0.045), which all had similar reductase activities (Figure 5b). In all strains, the culture supernatants displayed substantially higher iron-reducing capacities with no significant differences between strains (Figure S11b). These findings demonstrate that cell wall-bound melanin can reduce ferric iron, while extracellular components dominate iron reduction in the supernatant. Further, the iron content of the biomass – an indication of iron adsorption – was measured for the melanized WT and the non-melanized Δ*pks1* mutant after cultivation under different iron concentrations. Across all conditions, the Δ*pks1* mutant consistently showed significantly lower iron levels in its biomass compared to the WT: 45 % (*p* = 0.0039) on −Fe, 41 % (*p* = 0.0018) on +Fe, and 42 % (*p* = 0.0071) on hFe (Figure 5c).

To investigate whether the iron adsorbed to melanin could serve as an extracellular iron reservoir or facilitate iron sequestration from the environment, another drop assay was performed. WT and Δ*pks1* strains were precultured under −Fe, +Fe and hFe conditions. Subsequently, cell suspensions were spotted onto −Fe agar with and without 4.5 µM BPS. Both strains exhibited comparable growth regardless of the iron conditions during preculture.

Finally, to determine whether melanin formation influences growth in presence of iron chelators, sensitivity assays were performed across different genetic backgrounds (WT, Δ*nps1*, Δ*ftr1-fet1*) with BPS, and EDTA, with and without citrate supplementation (Figure 5e). The deletion of *pks1* did not seemingly alter the sensitivity of the strains to iron chelators, indicating that melanin does not play a detectable role in chelator resistance under the tested conditions.

## Discussion

Iron is an indispensable micronutrient for all living organisms, driving intense competition for its acquisition. This competition is particularly evident in parasitic interactions, where hosts actively restrict iron availability to inhibit the growth of fungal invaders. Saprobic fungi, on the other hand, acquire iron by digesting dead organic material and may protect their nutrient sources from competitors through efficient iron uptake, rapid growth, and the production of toxic secondary metabolites. In contrast, mutualistic relationships – such as those found in lichens – are characterized by cooperation, where different organisms harmonize their nutrient acquisition strategies.

We hypothesized that iron uptake by rock-inhabiting black fungi is unconventional, resulting in a high sensitivity to iron chelators. These organisms (i) are considered free-living, i.e. they do not rely on interaction partners, although they may exist within microbial communities (25), (ii) colonize oligotrophic substrates exposed to sunlight, which imposes UV radiation, temperature fluctuations, and desiccation stress – conditions that limit the growth of fast-growing saprobes, (iii) constitutively produce melanin, a pigment with known iron-adsorbing properties (26, 35), and (iv) may obtain iron from the weathering of iron-containing rock substrates (36). Overall, black fungi inhabiting such extreme environments are likely not exposed to immediate competition for iron, which may have enabled the evolution of simplified, and more energy-efficient iron acquisition and storage strategies.

To investigate this hypothesis, we first employed a bioinformatics approach, aiming to compare the potential iron acquisition strategies of selected Chaetothyriales black fungi – including pathogenic, saprobic, and rock-inhabiting species – based on their genomic repertoires. The results demonstrate that the closely related rock inhabitants *K. petricola* and *K. marmoricola*, both repeatedly isolated from marble surfaces (37, 38), possess a reduced set of iron acquisition genes. This gene set is characterized by a single RIA complex, absence of key siderophore biosynthesis enzymes (*sidA*, *sidD*), lack of a vacuolar iron transporter, and the presence of a ferritin, which may facilitate intracellular iron storage as in other eukaryotes (39, 40). In contrast, *K. obscura*, isolated, for example, from a cleanroom facility or a gasoline tank (41) contains a slightly altered gene set. These differences may reflect ecological adaptation and could contribute to future refinement of the genus *Knufia*. The closely related pathogenic species *A. fulminans* possesses a second RIA complex, additional SITs, and a putative vacuolar iron transporter. This pattern is consistent with those of pathogenic Herpotrichiellaceae species, which typically contain at least two RIA complexes, a higher number of SITs, the complete set for SIA (*sidA*, *sidC*, *sidD*), and multiple CCC-type transporters. Although non-pathogenic black fungi possess fewer putative SITs and FREs than their pathogenic relatives, the presence of multiple copies – combined with the siderophore specificity of these transporters and reductases (8) – suggests that they are equipped to utilize a range of siderophores, albeit a narrower spectrum than pathogenic species. The ability to utilize xenosiderophores, including those not originating from fungi, provides a competitive advantage by conserving metabolic energy (42–45). In addition, black fungi may retrieve iron from the degradation of self-produced or foreign heme via the heme oxygenase (HMX1). Uptake of xenosiderophores or heme may occur incidentally when these compounds are encountered in the environment; however, the likelihood of acquiring them from living or dying microorganisms is considerably higher within subaerial biofilms. Through this analysis, we identified distinct patterns of iron acquisition genes in pathogenic versus non-pathogenic black fungal species. These findings will benefit from the availability of additional genome sequences of black fungi from diverse habitats, which are expected to emerge from the community science project STRES (https://stresblackfungi.org/) (46). With a broader dataset and verified ecological or physiological traits, it may eventually become possible to predict the pathogenic/opportunistic potential of black fungi based on their iron acquisition gene profiles.

Experimentally, the functionality of RIA and SIA systems in *K. petricola* was demonstrated (Figure 6). Under laboratory conditions simulating the free-living lifestyle, *K. petricola* primarily acquires iron via the RIA system, encoded by *ftr1* and *fet1*, and secondarily through the SIA pathway involving *nps1*. Mutants lacking both *ftr1-fet1* and *nps1* are nearly non-viable, indicating that RIA and SIA are the only relevant mechanisms in *K. petricola* – at least under laboratory conditions. Cross-feeding assays with the Δ*nps1*/Δ*ftr1-fet1* mutant as test strain demonstrated the presence of a secreted and diffusible siderophore in *K. petricola*, corroborating previous findings from a chrome azurol S assay in this species (47). Biosynthesis of the siderophore requires the sole SidC-like NRPS, NPS1. However, we excluded ferricrocin as the identity of this siderophore. Therefore, future work should aim to elucidate the nature of this compound.

**Figure 6:**
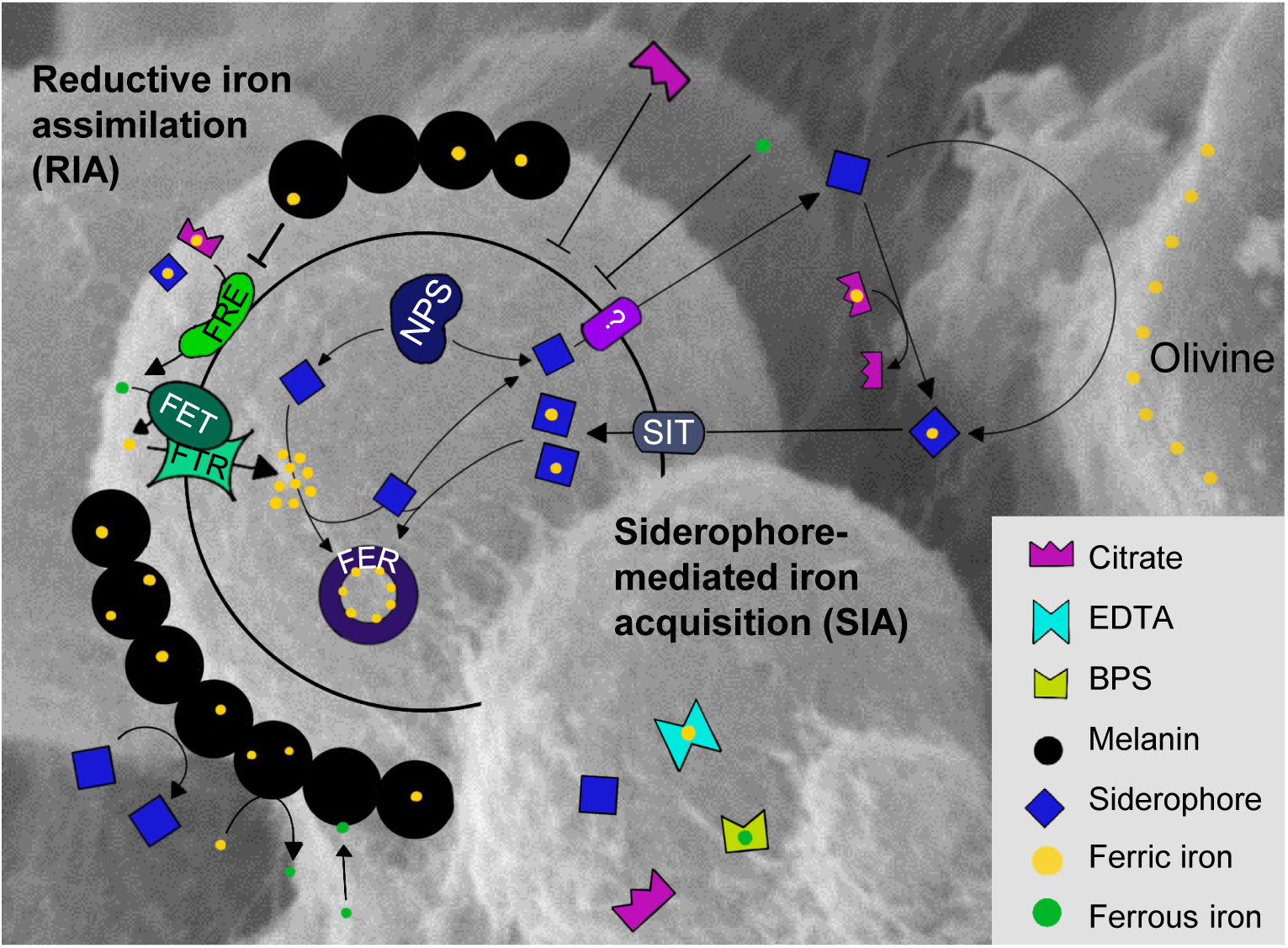
Proposed model of iron acquisition to intracellular storage in *K. petricola*. Iron can be taken up via a siderophore-mediated iron acquisition (SIA) system consisting of (xeno)siderophores and siderophore transporters (SIT1-4) but is primarily taken up via the reductive iron assimilation (RIA) system consisting of ferric reductases (FER1-12), a ferroxidase (FET1) and an iron permease (FTR1). The siderophore synthesized by the non-ribosomal peptide synthetase NPS1 remains unidentified and appears ineffective, making RIA the dominant mechanism and increasing susceptibility to strong iron chelators such as BPS and EDTA. Direct uptake of iron-bound citrate is not feasible; however, citrate-bound iron can be transferred to the RIA system or to the siderophore. Melanin, although adsorbing and reducing iron, does not contribute to iron uptake or storage as iron bound to melanin is not available for growth. Intracellular iron is likely stored in ferritin (FER1), as proteins for vacuolar iron storage are absent.

Iron is present at the surface of olivine either in the form of ferric (oxyhydr)oxide precipitates or as ferric iron within the structure of olivine (36). Whether the siderophore can accept iron from the former or the latter is unknown. Such a chemotropic response is, however, particularly remarkable, given that siderophore production was reduced in the presence of iron-rich olivine, suggesting that even low levels of the siderophore are sufficient to facilitate iron acquisition under these conditions. A lower siderophore production in the presence of an iron source was also observed for *A. fumigatus* (48) and *A. nidulans* (49), in which genes involved in both RIA and SIA are repressed by the GATA-transcription factor SreA under high iron while the bZIP transcription factor HapX represses iron-consuming pathways and activates siderophore production under iron limitation (50–52). Assuming a similar regulatory network for RIA and SIA genes in *K. petricola*, potentially governed by conserved transcriptional regulators, the disruption of RIA via Δ*ftr1-fet1* or SIA via Δ*nps1* may mimic iron-limiting conditions. This could, in turn, trigger compensatory upregulation of the remaining iron acquisition pathway, thereby enabling continued growth despite the loss of one system.

Addition of citrate alleviated iron limitation of *K. petricola* under iron-depleted conditions caused by chelators. However, the relatively high concentration required (i.e., 1 mM) reflects the low affinity of citrate – its binding constants are 10^9^ for ferrous iron and 10^12^ for ferric iron (53). The lack of improved growth of the Δ*nps1*/Δ*ftr1-fet1* mutant under iron-limiting conditions with citrate addition suggests that direct uptake of citrate-bound iron is unlikely. This aligns with the observation in *Ustilago sphaerogena,* where iron, but no citrate, was taken up when iron citrate was provided (54). The restored growth of both Δ*ftr1-fet1* and Δ*nps1* upon citrate addition, suggests that both the FREs of RIA and the siderophore of *K. petricola* can mobilize iron from citrate. The ability of the RIA system to utilize citrate-bound iron has also been demonstrated in *S. cerevisiae*, which lacks own siderophores (55) but can grow on ferric citrate as an iron source (56). Additionally, siderophores have been shown to acquire iron from ferric oxides via low-molecular weight acids like citrate (57, 58), supporting the idea of iron handover from citrate to siderophores.

We demonstrated that the cell wall-bound melanin of *K. petricola* is capable to adsorb and reduce iron. Iron adsorption capacity, inferred from the iron content of the biomass, correlated with iron availability in the culture medium for both the WT and the Δ*pks1* mutant. However, across all conditions, the non-melanized Δ*pks1* mutant consistently showed lower iron accumulation compared to the melanized WT, demonstrating that melanin adsorbs iron. This finding aligns with earlier studies reporting that melanized fungal biomass tends to accumulate more metals like iron than albino biomass, and that fungal melanin itself could adsorb metals (35, 59–61). Similarly, the higher iron-reducing capacity observed for the melanized biomass of *K. petricola* (WT) compared to the non-melanized biomass (Δ*pks1*) is consistent with previous reports on the iron-reducing properties of enzymatically produced DOPA (3,4-dihydroxyphenylalanine) melanin (62), autoxidized DOPA melanin (63), and DOPA melanin-producing *Cryptococcus neoformans* (64). This reduction is likely mediated by reduced catechol compounds of the melanin, which are hydrophilic and therefore positioned at the surface, facilitating interaction with iron ions (61). The observation that the higher iron content of the WT inoculants compared to that of the Δ*pks1* inoculants did not translate into reduced sensitivity towards BPS, suggests that the additional melanin-bound iron in the WT was not readily available for supporting the growth of new cells. Similarly and in contrast to other fungi – where disruption of iron uptake pathways typically results in both reduced cellular iron contents and impaired growth under low-iron conditions (17, 55, 65, 66) – the growth defect of the (melanized) Δ*ftr1-fet1* mutant on iron-limited media did not correlate with a decrease in total iron content. These findings further support the notion that iron adsorbed to the melanin cannot be mobilized to sustain growth. Considering that melanin can compete with chelators such as citrate, adenosine 5’-diphosphate (ADP) and possibly EDTA for both ferrous and ferric iron (61, 63), this lack of bioavailability is reasonable. The observation that melanized and non-melanized strains exhibited similar sensitivity to iron depletion caused by chelators suggests that melanin does not sequester iron in a way that significantly interferes with either RIA or SIA. Apparently, the kinetics of both high-affinity iron uptake systems outcompete the iron adsorption of melanin. Taken together, these observations indicate that, while melanin of *K. petricola* can adsorb and reduce iron, it neither impairs nor enhances iron acquisition via RIA or SIA. Instead, melanin appears to function as a passive iron sink, adsorbing lingering iron making it no longer accessible to both uptake systems – and unable to react with H_2_O_2_ to generate reactive oxygen species (ROS) via the Fenton reaction (67, 68). The ability of melanin to bind and stabilize iron in a non-reactive form likely contributes to cellular protection, providing an explanation why the non-melanized *K. petricola* strains were found more sensitive towards oxidative stress than their non-melanized counterparts.

The addition of chelators to the cultivation medium mimics the natural competition for chelated iron, where only organisms equipped with strong siderophores can mobilize iron from weaker chelators. In *K. petricola*, growth inhibition by the chelators BPS and EDTA in drop assays using iron-free MM began at concentrations as low as 4.5 µM and 4.0 µM, respectively. MICs, determined based on colony formation from plated CFU on MEA, were 150 µM for BPS and 200–250 µM for EDTA. These MIC values are markedly lower than those observed for *S. cerevisiae* under identical experimental conditions, indicating *K. petricola* and the other rock-inhabiting fungi are more sensitive to iron chelation than *S. cerevisiae*. This increased sensitivity implies that these fungi possess insufficient iron to sustain growth, potentially because of restricted iron acquisition and/or storage. To facilitate comparisons, we compiled MIC data for BPS and EDTA across various fungal species (Table S4), acknowledging that MIC values are often difficult to compare between studies due to variations in media composition, experimental setups, and particularly inoculum size – as highlighted by Kubo et al. (69). This overview reveals that *K. petricola* and related rock-inhabitants exhibit MICs for these strong iron chelators that are one to two orders of magnitude lower than those reported for other fungi.

The unusually low MICs of BPS and EDTA observed for rock-inhabiting fungi may result from a combination of factors. Among these, siderophore production stands out as the most compelling explanation: siderophore-deficient strains of *K. petricola* and other fungi (17, 70) consistently show higher sensitivity to iron chelators compared to their respective WTs. Despite the observed properties of the *K. petricola* siderophore *–* such as its secretion, diffusion through the agar, uptake by neighboring cells, and chemotropic behavior *–* it appears to be relatively ineffective in acquiring iron. Typically, siderophores bind ferric iron with very high affinity, exhibiting binding constants in the range of 10^25^ and 10^62^ (71). For comparison, BPS binds ferrous iron with a binding constant of 10^21^ (72), while EDTA binds ferric iron with a binding constant of 10^25^ (73). In addition to the possibility that the *K. petricola* siderophore has a relatively low affinity for iron, its limited effectiveness may also be due to low production levels or a predominant localization in the cytosol or at the plasma membrane. Especially in oligotrophic environments, an amphiphilic siderophore that remains predominantly associated with the plasma membrane and only occasionally diffuses into the extracellular space may offer a distinct advantage (74). By retaining the siderophore at the cell surface, the rock-inhabiting fungi may reduce its loss to the surroundings and thereby save metabolic energy that would otherwise be lost by continuous siderophore synthesis. To date, amphiphilic (coprogen) siderophores have been reported from one fungal species, *Trichoderma hypoxylon* (75).

Siderophore production does not necessarily correlate with higher MIC values for EDTA and BPS. For example, the yeasts *Candida* species and *S. cerevisiae* exhibit relatively high MICs despite their inability to produce siderophores – values that exceed those observed for the tested black fungi (Table S4). However, melanin production as shared trait among the four rock-inhabiting fungi is not the underlying reason. Instead, the relatively high MIC values in *S. cerevisiae*, indicating higher tolerance to iron deprivation, may be explained by efficient iron storage, particularly the sequestration of iron in vacuoles mediated through CCC1 (76) and its remobilization by an additional RIA (FET5–FTH1) complex (77). As genes for vacuolar iron storage are absent from *K. petricola* and *K. marmoricola*, it is plausible that their limited tolerance to iron-depleted conditions (reflected in low MIC values) result also from insufficient intracellular iron storage. This deficiency may not be sufficiently compensated by the ferritin, assuming the gene is expressed and produces a functional iron-binding protein.

Irrespective of the underlying reason, the low MICs of *K. petricola* and other rock-inhabiting fungi open avenues for the use of strong iron chelators in a mitigation strategy to prevent material colonization. These fungi not only colonize stone surfaces of culturally significant monuments (78) but also energy-generating photovoltaic cells (27, 79, 80). Together with other organisms and abiogenic deposits, rock-inhabiting fungi block solar radiation, having been shown to lower the efficiency of the solar panels by up to 11 % after 18 months (81). Removing or preventing the formation of the typical microcolonies of these fungi has proven challenging (29). Considering the urgent need to derive more fossil fuel-free energy (82), a novel approach using biocides would therefore be most welcome. EDTA, although not considered fungicidal or only inhibitory at higher concentrations (83, 84), prevented growth of rock-inhabiting species at relatively low concentrations (i.e., acts fungistatic). The EU Risk Assessment Report for EDTA (85) states that one should limit the risks due to the high emissions in an industrial context, leading to risks for aquatic organisms. The application of EDTA on plant leaves as a fertilizer (assumed the worst-case approach) is not considered a risk for terrestrial organisms (85), even though the chelated form is commonly applied with concentrations as high as 2.5 mM (86). Nevertheless, care should be taken to use chelators which are more easily degradable than EDTA such as N,N’-ethylenediaminedisuccinic acid (EDDS) (87–89), iminodisuccinic acid (IDS) (88, 89) or 2-((1,2-dicarboxyethyl)amino)pentanedioic acid (90). To exert a fungistatic effect, these chelators still need to be sufficiently strong. IDS has a binding constant for ferric iron of 10^14^ (91), only marginally higher than the binding constant of citrate for ferric iron of 10^12^ (53), and will thus not be useful. EDDS, however, has a binding constant of 10^22^ (92), comparable to EDTA (K = 10^25^) (73) and is therefore of interest within this context. Future work should focus on evaluating chelator-based approaches under real-world conditions to develop sustainable strategies for preventing fungal growth on exposed surfaces.

## Materials and methods

### Fungal culturing and reagents

*K. petricola* strain A95, isolated from a marble stone surface near the Philopappos monument in Athens, Greece (93), was used as wild type (WT) (Table S3). Other rock-inhabiting black fungi i.e., BAM-BF001 (Eurotiomycetes), BAM-BF027 (Arthoniomycetes), and BAM-BF046 (Dothideomycetes) were isolated from solar panels in Berlin, Germany (unpublished). *S. cerevisiae* strains used were DSM 1333 and FY843 (94). Fungal cultures were maintained on malt extract agar (MEA), containing 20.0 g l^-1^ glucose, 0.1 g l^-1^ casein peptone, 20.0 g l^-1^ malt extract, and 20.0 g l ^1^ Kobe agar. Unless stated otherwise, experiments were conducted with a minimal medium (MM), containing 701 mM D-sucrose, 35.3 mM NaNO_3_, 5.878 mM H_2_KPO_4_, 12.59 mM KCl, 2.029 mM MgSO_4_*7H_2_O, 10.0 µM CaCl_2_*2H_2_O, 1.002 µM ZnSO_4_*7H_2_O, 1.001 µM MnCl_2_, 1.000 µM CuSO_4_*5H_2_O, 1.00 µM H_3_BO_3_, 0.101 µM CoCl_2_*6H_2_O, 0.0992 µM Na_2_MoO_4_*2H_2_O, 0.023 µM Na_2_SeO_3_*5H_2_O, 0.101 µM NiCl_2_*6H_2_O and 15.0 g l^-1^ bacteriology grade agar, either adding no (-Fe), 30 µM (+Fe) or 1,511 µM (hFe) FeSO_4_*7H_2_O. All glassware was cleaned overnight with 1 M HNO_3_ to prevent any chemical contamination. Media were buffered at pH 6 with 12.30 mM 2-(N-morpholino)ethanesulfonic acid (MES), added after autoclaving. Cultures were incubated in the dark at 25 °C. Biomass of all strains was disaggregated by glass beads (3-4 mm diameter) in a Retsch mixer mill (10 min at 30 Hz), and washed twice with sterile MilliQ water. Numbers of colony forming units (CFU), i.e. single cells and cell aggregates, were determined using a Thoma cell counting chamber.

### Growth assays

All strains were precultured for seven days on –Fe agar, except the Δ*nps1*/Δ*ftr1-fet1* mutant which was precultured on hFe agar to promote growth. Δ*nps1* mutants were never grown in the same Petri dish as *nps1*-expressing strains to avoid cross-feeding by other strains. Growth on different solid media was tested by semi-quantitative drop assays. For this, 10 µl of serial dilutions with 10^6^, 10^5^, 10^4^, and (in certain cases) 10^3^ CFU ml^-1^ were dropped onto solid medium with or without Fe. Varying concentrations of iron chelators (bathophenanthroline disulfonate (BPS), Na_2_EDTA*2H_2_O or citrate) were added after autoclaving. For inducing oxidative stress, H_2_O_2_ was added to the medium after autoclaving. To test the use of melanin as an iron source, WT and Δ*pks1* were grown for seven days on either –Fe, +Fe or hFe agar covered with cellophane, harvested and treated as described before, and point-inoculated on –Fe with and without 4.5 µM BPS. For the cross-feeding assay, –Fe agar, without additive or supplemented with 2.5 mg ml ^1^ synthetic forsterite or 2.5 mg ml^-1^ natural iron-containing olivine (India, Hausen Mineraliengrosshandel both ground to < 63 µm) was inoculated with four 10 µl drops containing 10^3^ CFU of Δ*nps1*/Δ*ftr1-fet1* in between of which (and with a distance of 20 mm) one 10 µl drop (10^3^ CFU) of the test strain was dropped. To test whether strains can grow towards an iron source, 50 mg of the same ground olivine and forsterite were streaked out on opposite sides of rectangular Petri dishes containing –Fe agar. Exactly in between the mineral streaks, at a distance of 50 mm, 10 µl drops containing 10^3^ CFU of the test strains were spotted in a row. After 74 days, images were taken and the distances of growth from the initial drop towards olivine and forsterite were quantified. The ratio of growth towards olivine vs. towards forsterite (i.e., the relative growth towards olivine) was calculated for all five colonies of each Petri dish and the average was taken. Each growth assay was conducted at least three times with reproducible outcomes. Representative images from a single experiment are presented.

The lag phases and growth rates of individual colonies was determined using the ScanLag system (95). For this, –Fe, +Fe and hFe agar was inoculated with 500 CFU of the respective strains. The Petri dishes (9 cm in diameter) were subsequently incubated in flatbed scanners (V600, Epson, connected to a computer running Windows 10 OS) in the dark at 25 °C. Scans were taken every six hours using a custom PowerShell script. The lag phase and growth rate were quantified using a custom Matlab script based on the Matlab code of I. Levin-Reisman et al. (95), assuming linear growth. As a slight inhibition effect was observed (i.e., a decrease in the growth rate with an increasing CFU, data not shown), only Petri dishes with a CFU in the range of 50 to 200 (except for Δ*nps1*/Δ*ftr1-fet1* which had CFUs in the range of 6 to 23, which might have caused an overestimation of its growth rate) were analyzed.

To define the minimum inhibitory concentration (MIC) of EDTA and BPS, 20 ml MEA per Petri dish (9 cm), buffered at pH 6 with 12.30 mM MES and supplemented with different concentrations of EDTA or BPS, was inoculated with 200 to 450 CFU (three independent replicates were run per condition) and incubated for 14 days in the dark. Relative colony growth in percent was calculated.

### Bioinformatics analyses

Analyses were performed using Geneious Prime 2024.0.7 (Biomatters Ltd.). Genome assemblies of selected species were retrieved from GenBank (Table S1) or were accessed from in-house resources (*K. petricola* A95 assembly v1, unpublished). *De novo* gene prediction and revision of incorrect gene models was performed using the Augustus plugin 0.1.1 (96) Candidate proteins were identified by internal BLAST searches (default parameters) using sequences from *S. cerevisiae* and *A. fumigatus* as queries. Conserved protein domains were identified using the InterProScan plugin 2.1.0 (97). Sequences of the *K. petricola* iron acquisition genes have been submitted to GenBank (Table S2). All protein sequences used for analyses in Figure 1 are listed in Table S1. Protein sequences were aligned using MUSCLE v5.1 with default parameters. Phylogenetic trees were inferred with the PHYML plugin 2.2.4 (98), applying the LG amino acid substitution model and 100 bootstrap replicates. CRISPR/Cas9 target sites (protospacer with protospacer-adjacent motif ‘NGG’) in the genomic loci of interest were identified using the CRISPR Finder tool (default settings), with the *K. petricola* genome sequence as the off-target reference database. The protospacers used had at least four mismatches to the closest predicted off-target site. SnapGene 4.0.8 (GSL Biotech LLC) was used to design and document cloning strategies by generating sequence maps and aligning Sanger sequencing reads.

### Standard molecular methods

Genomic DNA was extracted as described by Voigt, Knabe et al. (30). DNA was mixed with Midori Green Direct (Biozym Scientific) and separated in 1 % agarose gels with the MassRuler DNA Ladder Mix (Thermo Scientific) or the 1 kb Plus DNA Ladder (New England Biolabs, NEB) as a size reference. PCR reactions were carried out with primers from Eurofins Genomics (Table S5), the Q5 High-Fidelity DNA polymerase (NEB) for cloning and sequencing and the *Taq* DNA polymerase (NEB) for diagnostic PCR. Plasmids listed in Table S6 were assembled via homologous recombination in *S. cerevisiae* (99, 100). Plasmids were amplified in *Escherichia coli* DH5α (Invitrogen) and isolated from *S. cerevisiae* and *E. coli* with the Monarch Plasmid Miniprep Kit (NEB). Larger amounts of plasmid were extracted from *E. coli* with the NucleoBond Xtra Midi Kit (Macherey-Nagel). PCR products were purified with the Monarch PCR and DNA Cleanup Kit (NEB) and sequenced with the Mix2Seq Kit at Eurofins Genomics. sgRNAs were synthesized *in vitro* with the EnGen sgRNA Synthesis Kit (NEB), purified with the Monarch RNA Cleanup Kit (NEB), and assembled in a molar ratio of 1:1 with EnGen Spy Cas9 NLS (NEB) by incubation for 10 min at 22 °C.

### *Genetic manipulation of* K. petricola

The deletion of coding regions (knock-out, KO) and the targeted integration of expression constructs (knock-in, KI) into the *K. petricola* genome was accomplished by introducing double strand breaks (DSB) by the CRISPR/Cas9 technology and providing donor DNA for repair of the DSB by homologous recombination (HR) according to Voigt, Knabe et al. (30) and E. A. Erdmann et al. (31). Target-specific ribonucleoproteins (RNP) were either expressed and assembled *in-vivo* from plasmid DNA or added along with the donor DNA as pre-assembled RNP as specified in Table S7. Per approach, 1 x 10^6^ protoplasts were transformed with 10 to 15 µl of a PCR sample or a restriction digest (linear donor DNA) together with either 2 µg of sgRNA- and Cas9-delivering plasmid (circular DNA), or 1 µg of sgRNA assembled with 5 µg of Cas9, as described previously. The correct integration of deletion and expression constructs as consequence of HR was detected by diagnostic PCR by combining primers binding upstream or downstream of the integration site with those binding within the integrated sequences i.e., in resistance or expression cassettes (Figure S6, Figure S8, Figure S10). For all genetic manipulation strategies, multiple independent transformants were generated and subjected to an initial phenotypic screening; representative data from one randomly selected transformant are presented.

### Determination of iron content in biomass

Solid MM –Fe, +Fe and hFe medium covered with cellophane was inoculated with 50,000 cells. After 21 days, the biomass in the center of the Petri dish was taken off and dried at 65 °C. A weighed amount was digested in polytetrafluorethylene beakers with 1 ml of 9.8 M H_2_O_2_ and 1 ml of 14.3 M HNO_3_ at 150 °C. After digestion, the acidic solution was evaporated, and the resulting pellet was dissolved in 4 ml of 1 M HNO_3_ via ultrasonication. The HNO_3_ solution containing the dissolved biomass residue was analyzed by inductively coupled plasma-optical emission spectroscopy (ICP-OES, Varian 720-ES) in the HELGES laboratory at GFZ, Potsdam (101) as described before in (36) (see Method S1, Table S8, Table S9 for details). The analytical uncertainty used to interpret the sample results is quantified based on the accuracy and precision of the repeated measurement of quality control standards, the error of the analysis of the standards, and the contribution of the blank (i.e., the solution used to dilute the samples, see supplementary information). The iron content was quantified by dividing the amount of iron in the digest by the amount of biomass digested (µg g^-1^).

### Iron reduction assay

To obtain the ferric reduction potential of the biomass, the absorbance of the red BPS-Fe(II) complex was measured spectrophotometrically. 20 ml of liquid –Fe medium in Erlenmeyer flasks were inoculated with 10^6^ CFU and placed on a shaker for 14 days in the dark at 25 °C and 100 rpm. Afterwards, the cultures were centrifuged, a sample of the supernatant was taken, and the pellet (biomass) was washed once with fresh –Fe medium. Fresh –Fe medium with 0.5 mM Fe(III)Cl_2_*6H_2_O and 0.5 mM BPS was added to the biomass and supernatant samples and incubated for 3 h in the dark at 25 °C and 100 rpm. After another centrifugation step, the A_535_ was measured using a UV-vis spectrophotometer (Genesys 10S, Thermo Scientific). An abiotic control was used as a blank. The pellet was dried at 65 °C for one day and the dry weight was measured. The concentration of reduced iron was obtained from the measured A_535_ values by running the experiment with a standard series (0, 0.01, 0.025, 0.05, 0.1, 0.25, 0.5 and 1 mM Fe(II)SO_4_*7H_2_O) of reduced iron (Figure S11a). Finally, the reduced iron concentration was divided by the dry weight and the duration of the incubation to obtain the reduction capacity in mM Fe(II) g^-1^ h^-1^. In the case of the supernatant samples, the reduced iron concentrations were divided by the dry weight of the respective culture, the fraction of supernatant analyzed and the duration of the incubation.

### Statistical analyses

In general, all data are shown as the average of three or four independent replicates with two times the standard error (Table S10). Such a sample size is too low to test for normality. Differences between the strains and conditions are thus analyzed via the non-parametric Kruskal Wallis test followed by the Conover-Iman post-hoc test with Benjamini-Hochberg p-value adjustment (alpha level = 0.05). For both the conover.test package (v1.1.6) in R software (v4.5.1; R core Team, (102)) was used.

## Data availability

Nucleotide and protein sequences of studied *K. petricola* genes are available from GenBank (https://www.ncbi.nlm.nih.gov/genbank/). *K. petricola* strains, generated in this study, are available upon request.

## Acknowledgment

We thank Niclas Nordholt for help with setting up the ScanLag system and Steffen Ganschow for providing us with forsterite. Eileen Erdmann, Sarah Nitsche, Jenny Straßner and Oliver Voigt are thanked for their help with cloning. Pedro Maria Martin-Sanchez and Christopher Gebhardt are thanked for the isolation of the rock-inhabiting fungi from solar panels. We are grateful for access to the ICP-OES of the HELGES laboratory. This study was funded by internal funds of the BAM.

## Author contributions

**RG**: Conceptualization, Data Curation, Formal Analysis, Investigation, Methodology, Resources, Validation, Visualization, Writing – Original Draft, Review & Editing. **JS**: Conceptualization, Data Curation, Formal Analysis, Investigation, Supervision, Writing – Original Draft, Review & Editing. **AAG**: Conceptualization, Funding Acquisition, Supervision, Writing – Review & Editing.

